# CellNetVis: a web tool for visualization of biological networks using force-directed layout constrained by cellular components

**DOI:** 10.1101/163410

**Authors:** Henry Heberle, Marcelo Falsarella Carazzolle, Guilherme P. Telles, Gabriela Vaz Meirelles, Rosane Minghim

## Abstract

**Background:** The advent of “omics” science has brought new perspectives in contemporary biology through the high-throughput analyses of molecular interactions, providing new clues in protein/gene function and in the organization of biological pathways. Biomolecular interaction networks, or graphs, are simple abstract representations where the components of a cell (e.g. proteins, metabolites etc.) are represented by nodes and their interactions are represented by edges. An appropriate visualization of data is crucial for understanding such networks, since pathways are related to functions that occur in specific regions of the cell. The force-directed layout is an important and widely used technique to draw networks according to their topologies. Placing the networks into cellular compartments helps to quickly identify where network elements are located and, more specifically, concentrated. Currently, only a few tools provide the capability of visually organizing networks by cellular compartments. Most of them cannot handle large and dense networks. Even for small networks with hundreds of nodes the available tools are not able to reposition the network while the user is interacting, limiting the visual exploration capability.

**Results:** Here we propose CellNetVis, a web tool to easily display biological networks in a cell diagram employing a constrained force-directed layout algorithm. The tool is freely available and open-source. It was originally designed for networks generated by the Integrated Interactome System and can be used with networks from others databases, like InnateDB.

**Conclusions:** CellNetVis has demonstrated to be applicable for dynamic investigation of complex networks over a consistent representation of a cell on the Web, with capabilities not matched elsewhere.

## Background

With the advent of “omics” science, analyses performed from screening a wide range of physical, genetic and chemical-genetic interactions have brought new perspectives to contemporary biology, as they provide new clues in protein/gene function, help to understand how metabolic, regulatory and signaling pathways are organized and facilitate the validation of therapeutic targets and potential drugs. Biomolecular interaction networks are simple abstract representations where the components of a cell (e.g. genes, proteins, metabolites, miRNAs etc.) are represented by nodes and their inter-actions are represented by edges. An appropriate display of the data is crucial for understanding such networks, particularly regarding high-throughput analysis.

Since different regions of the cell are related to specific activities, visually organizing network nodes into cellular components can help understand the biological system and its relationship to the distribution of network elements over the cell structure. The position of nodes can unveil, for instance, patterns of relations among different cellular components. Additionally, it is common to query just a subnetwork of an entire interactome, so when users query specific pathways by a list of their units (e.g. gene symbols) they can easily see, by using a proper layout, where these pathways may occur in the cell.

Many tools are available to visualize and explore network models but most of them are not designed to partition networks into a cell structure. Among those are Graphviz [1], Gephi [2], Pajek [3], PEx-Graph [4], Cystocape [5] and Tulip [6]. They were created for a generic purpose, being applied in problems ranging from social network analysis to biology. Cytoscape is the most popular tool in biology and counts with many plugins for Systems Biology in particular, including two that work with cellular partitions: Cerebral [7] and Mosaic [8]. Other software systems, like Extended LineSets [9], Entourage [10], and ReactionFlow [11], focus on the analysis of pathways and their mechanisms.

Garcia et al. describe an extension to the force-directed layout to place nodes according to their connection and class structure [12]. In their method, the cellular component annotations can define the class structure and approximate nodes of the same class. The approach, however, does not represent cellular components. Other approaches that group nodes in two-dimensional space have been proposed, such as constrained force-directed layout [13], constrained projections [14], hierarchical graph placement [15, 16, 17] and others [18, 19, 20, 21]. Despite their good performance even for large networks, the cell structure is not taken into consideration in either of those cases. Also, they are not adapted to display networks in an explicitly full cell diagram.

Only a few tools provide the capability of displaying networks organized by cellular components. Biographer [22] is a web-based tool to edit and render reaction networks. It implements features for visualization based on Systems Biology Graphical Notations (SBGN). The user can manually create shapes of type “compartment” and position nodes inside them. Mosaic [8] is a Cytoscape plugin and can represent a network divided into cellular partitions automatically, duplicating nodes when there is more than one cellular component annotation. It uses force-directed layout, but it does not update the layout when nodes are moved. Also, the display was designed to show small subnetworks. Cerebral [7, 23], originally designed as a Cytoscape [5] plugin and extended to work with Cytoscape.js, can automatically divide the network into subcellular regions represented by parallel rectangles, one over the other, which is not consistent with the standard graphical representation of a cell. Kojima et al. developed a grid layout that may be applied over a full cell diagram, representing the cellular components properly [24]. The new version, Cell Illustrator Online [25], is a tool that enables drawing, visualization and modeling of biological pathways. It produces layouts that more closely resemble a consistent cell diagram and displays a network across cellular components. However, that tool is more focused on the mechanisms rather than on the network overview and exploration, the structure is manually defined by the user, and it is neither free nor open-source.

Despite the capability of drawing networks organized by cellular components, Mosaic [8], Cerebral [7], CerebralWeb [23] and Cell Illustrator [24] do not provide real-time automatic layout modifications for dynamic exploration. Even for small networks, with hundreds of nodes, these tools cannot reposition the network while the user is interacting, exploring the layout and manually repositioning the nodes. Many biological networks are dense causing the “hairball” problem, what makes the analysis of links, flows and topology difficult. Interactively moving nodes or organelles can increase readability and understanding, clarifying the flow of edges between them and letting the user explore the view to better understand the network dynamics.

We have developed a web tool called CellNetVis that tackles most of the mentioned drawbacks. It is meant for easy and dynamic display and exploration of biological networks over a full cell diagram. It uses an iterative force-directed algorithm to produce a dynamic layout for the entire network where nodes are positioned into movable cellular components. The input for the tool is a properly annotated network in the XGMML format. The tool displays the network over a standard cell graphical representation showing the main partitions and organelles according to the Gene Ontology (GO) cellular component database [26]. It also provides interactive features such as search, selection, drag and drop of organelles and nodes, as well as the capability of displaying nodes annotation information.

CellNetVis allows certain features, essential to current biological network analysis needs, not provided by other tools, such as, at the same time, being webbased, supporting large networks and providing automatic display of nodes inside their cellular components. Additionally, the particular implementation of the force-directed algorithm provides a balance between processing time and visual understanding of network structure with layout flexible to adapt to user’s manipulation. We discuss these issues in contrast with available tools in the section titled *Comparison with available tools*.

## Implementation

CellNetVis was written in Javascript and HTML and is a free and open-source software. It loads networks constructed using the XGMML format [27]. The only requirement is that the network nodes must have an attribute named either “Selected CC” or “Localization”, which corresponds to a unique selected cellular component (CC), such as the one generated by the IIS [28] and the InnateDB [29]. As the majority of proteins are described as acting in more than one subcellular compartment in GO, IIS and InnateDB apply a priority filter to assign the most specific cellular component to each protein, which is then used by CellNetVis to position the nodes in the cell diagram. Other strategies for assigning a single cellular component to each node may be adopted as well.

As shown in Table 1, InnateDB specifies in the XGMML file five possible compartments, while the IIS specifies twenty-one. CellNetVis works with all these 21 compartments. Additionally, the tool supports the retrieval of cellular components for human, mouse and bovine genes from the InnateDB web service. In this case, nodes must have an attribute that identifies the gene or protein ID in the Ensembl [30], Entrez [31], InnateDB [29] or UniProt [32] format.

**Table 1:** Cellular components specified in the XGMML file by IIS and InnateDB.

| IIS | InnateDB |
| --- | --- |
| Extracellular, cell wall, plasma membrane, mitochondrion, endoplasmic reticulum, Golgi apparatus, endosome, centrosome, microtubule organizing center, lysosome, vacuole, glyoxysome, glycosome, peroxisome, amyloplast, apicoplast, chloroplast, plastid, cytoplasm, cytosol, and nucleus. | extracellular, cell surface, plasma membrane, cytoplasm, and nucleus. |

A few decisions guided the construction of the cellular design in CellNetVis. Figure 1 shows an example of a small network displayed over the cell diagram. The cell is drawn aiming to highlight the separation between the main subcellular compartments: extracellular region, cell wall, plasma membrane, cytoplasm, and nucleus. Cell contour lines are drawn using lighter colors as they serve only as a reference. In contrast, network nodes contour lines are displayed with darker colors by default, and if nodes are selected, then the remaining ones are shown with transparency to improve contrast. Regarding the organelles, their contour lines are drawn with less contrast to reduce visual density, since typically these are regions with many edge crossings. The cell diagram is colored using a ColorBrewer [33] “BrBG” diverging scheme, characterized by colors that can be easily differentiated.

**Figure 1:**
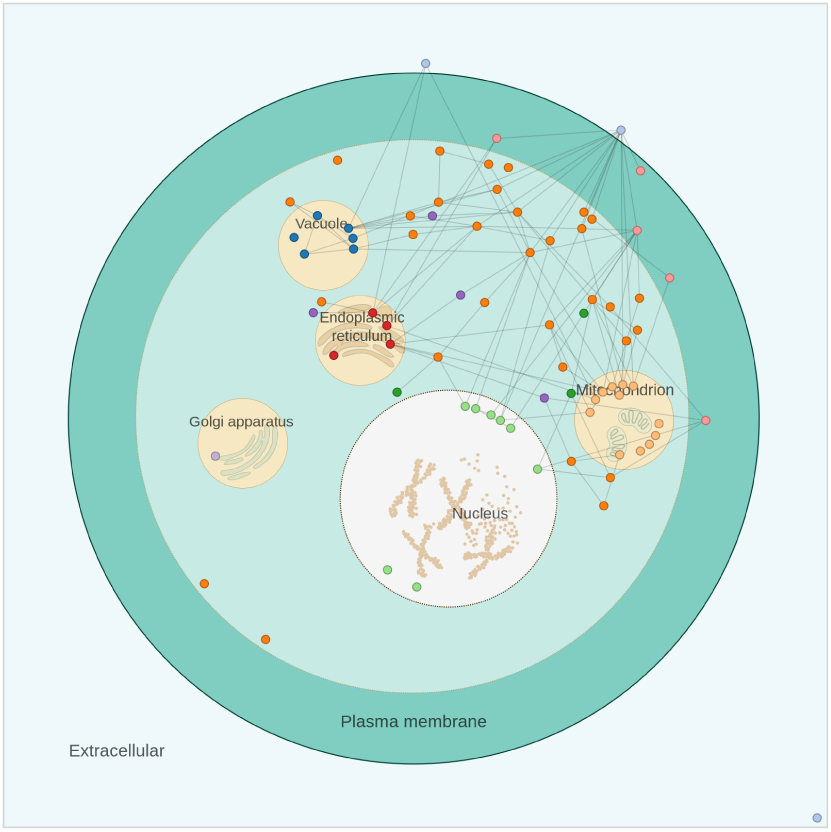
Display of a small network over a cell diagram.

If the compartment attribute is annotated with any other value not specified on CellNetVis or is empty, it will be positioned in the cytosol. If a node is also annotated with a “Cellular Component” multivalued attribute, all compartments in the list will be highlighted when the node is selected. For instance, if the protein A is drawn on *nucleus* (Selected CC) and has “nucleus, cytosol, mitochondrion” annotated in the “Cellular Component” attribute, all these components will be highlighted when the user selects this vertex. The user can also change the value of “Selected CC” during the visualization process.

The network is drawn over the cell representation by the force-directed layout adapted from the algorithm implemented in D3 [34] version 3.0 (https://github.com/d3/d3/wiki/Force-Layout). Our layout has the important advantage over existing tools based on grid-layout of enabling dynamic plotting and interaction with complex networks. We have modified the force-directed layout to constrain the movement of each node to the area of its respective cellular component.

Since the constraints computation in the force-directed layout is computationally expensive, the cell diagram is drawn using only circles, instead of other shapes that are commonly used to create a cell diagram. Complex shapes increase the time to check if each node is in the correct region and, given its current position, recalculate the new position according to the respective component shape. Circles simplify these verifications and position calculations. Another thing that reduces calculations is allowing movement of organelles and their content to extracellular region. The control over the cell structure consistency is left to the user’s discretion.

During each iteration of the force-directed algorithm, the position *x, y* of each node *n* is updated. How *x, y* is recalculated depends on the Selected CC (*n.cc*) of *n*, as described in the pseudo-code below.

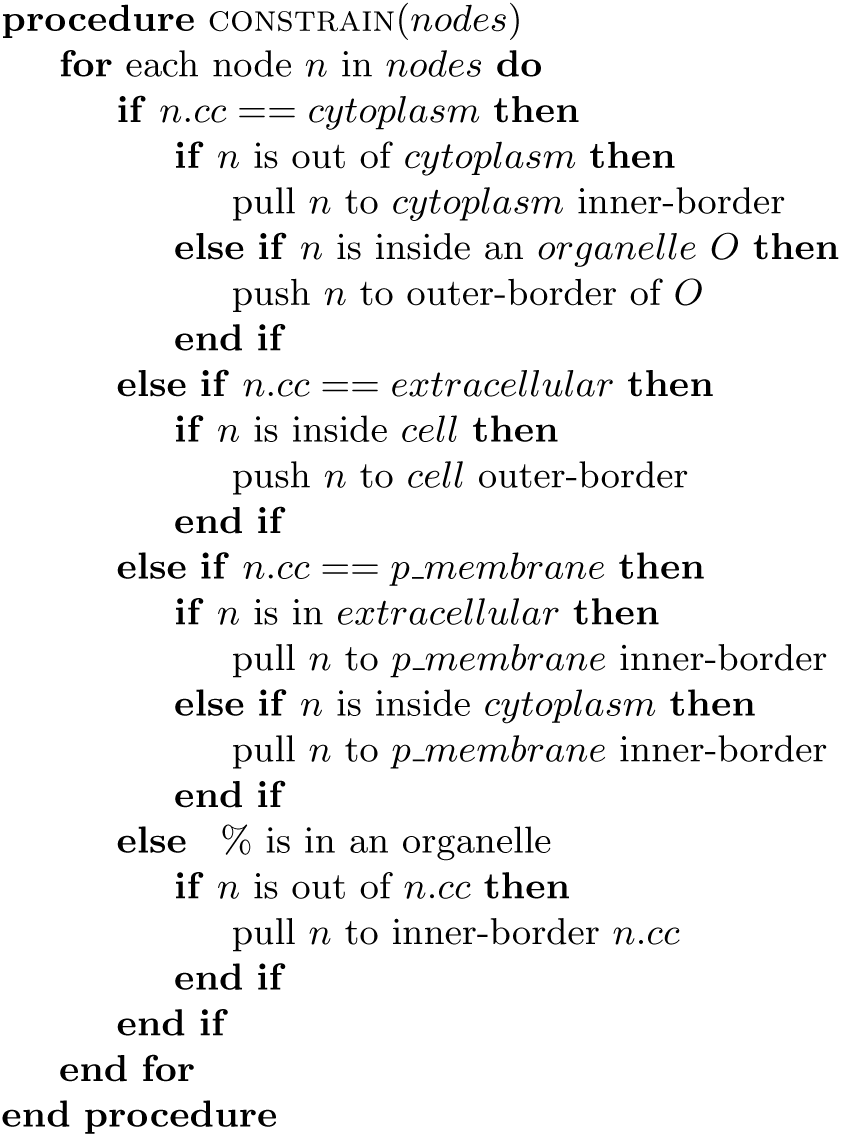

When the node is in an organelle, the algorithm checks the distance *R* between the center of a node (point *A*: *x, y*) and the center of its corresponding cellular compartment’s circle (point *B*: *cx, cy*) (Additional File 1A). The node is then placed in the new position, point A’ (*x*^*i*^, *y*^*i*^) calculated by 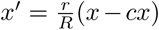 and 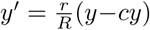, where 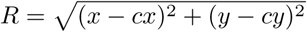 and *r* is the organelle radius. When the node is in the cytosol, the computation is similar, but in the opposite direction (Additional File 1B). When the node is in the cell wall or in the plasma membrane, two constraints are checked since there is an outer limit (cell wall or extracellular regions) and an inner limit (cytoplasm or plasma membrane).

When nodes are constrained to specific cellular regions, edges cross at higher rates in the layout. If the network is large, there will probably be too many nodes in the organelles and forces pulling nodes in the same region of the compartment, resulting in overlap of nodes. This limitation is not a feature of CellNetVis, but a deep problem in graph drawing. To reduce this effect, a new constraint was implemented in CellNetVis. The algorithm identifies whether a node is colliding with another one. If so, the nodes are repositioned. This verification is done after each iteration of the D3 force placement and queries a quad-tree data structure [35].

A user controlled parameter, named *repulsive* is used to support overlap reduction procedures. If *repulsive* is large, layout stability is lower but the visual separation of nodes is faster. If *repulsive* is small, visual stability is higher, but the nodes need more time to separate. Smaller *repulsive* values do not guarantee that nodes will not overlap. When a large network is loaded, this procedure is disabled by default. Other parameters of the force-directed algorithm can be configured. For instance, setting the *charge* of each node to a more negative value will make nodes more separated. All parameters available in CellNetVis are further explained in the Help page.

The user has four additional options to improve the network layout: moving organelles, constraining nodes to a specific position, hiding unfocused nodes (filter function) and turning on edge bundling, that is used to decrease cluttering from crossing edges. Organelles that are not annotated in any node of the network are removed from the view. We integrated the Corneliu Sugar implementation (https://github.com/upphiminn/d3.ForceBundle) of ForceDirected Edge Bundling [36] in CellNetVis.

Highlighting neighborhoods of selected nodes, displaying labels, calculating network topology measures and the possibility to color nodes according to different attributes were implemented. Counting of nodes per cellular component was also implemented as a donut chart. The cell diagrams can be exported as a bitmap (.PNG) or vector (.SVG) image.

To allow integration with other systems and publication of a network view in the form of a URL, CellNetVis provides a special parameter named “file”, which receives the URL of a XGMML file. When this parameter is used, an asynchronous call is executed by CellNetVis and, after the successful download, the file is parsed and processed the same way as a regular input. The external XGMML server provider must have the CORS header ‘Access-Control-AllowOrigin’ set (https://www.w3.org/TR/cors/#access-control-allow-origin-response-header).

The response time of CellNetVis depends on the time taken for the construction of the network structure by the Javascript code, the SVG rendering time taken by the web browser and, if the URL approach is used to load the XGMML file, the time to download the network. All the computation is done on the client-side, so the time needed to display a network and interact with the system depends only on the user’s computer.

## Results and discussion

CellNetVis is capable of displaying information related to complex networks, nodes, and edges as well as their relations with cell partitions. Figure 2 shows the CellNetVis interface. To analyze a network in the cell diagram, the user starts by uploading a network as a XGMML file (Figure 2A). The network will be loaded in the cell diagram area (Figure 2G) and the nodes will be distributed inside each subcellular localization according to its annotation. Alternatively, the user may create an URL that specifies the “file” parameter, that is, the CellNetVis URL plus the XGMML file URL. The force-directed algorithm starts automatically when a network is loaded and will resolve the positioning of nodes within each cellular component. It may be interrupted and restarted at any time (Figure 2D). Nodes and organelles may be manually positioned along the display. When that is done, the neighboring nodes or nodes inside the moving cellular components will be moved accordingly (Figure 2G).

**Figure 2:**
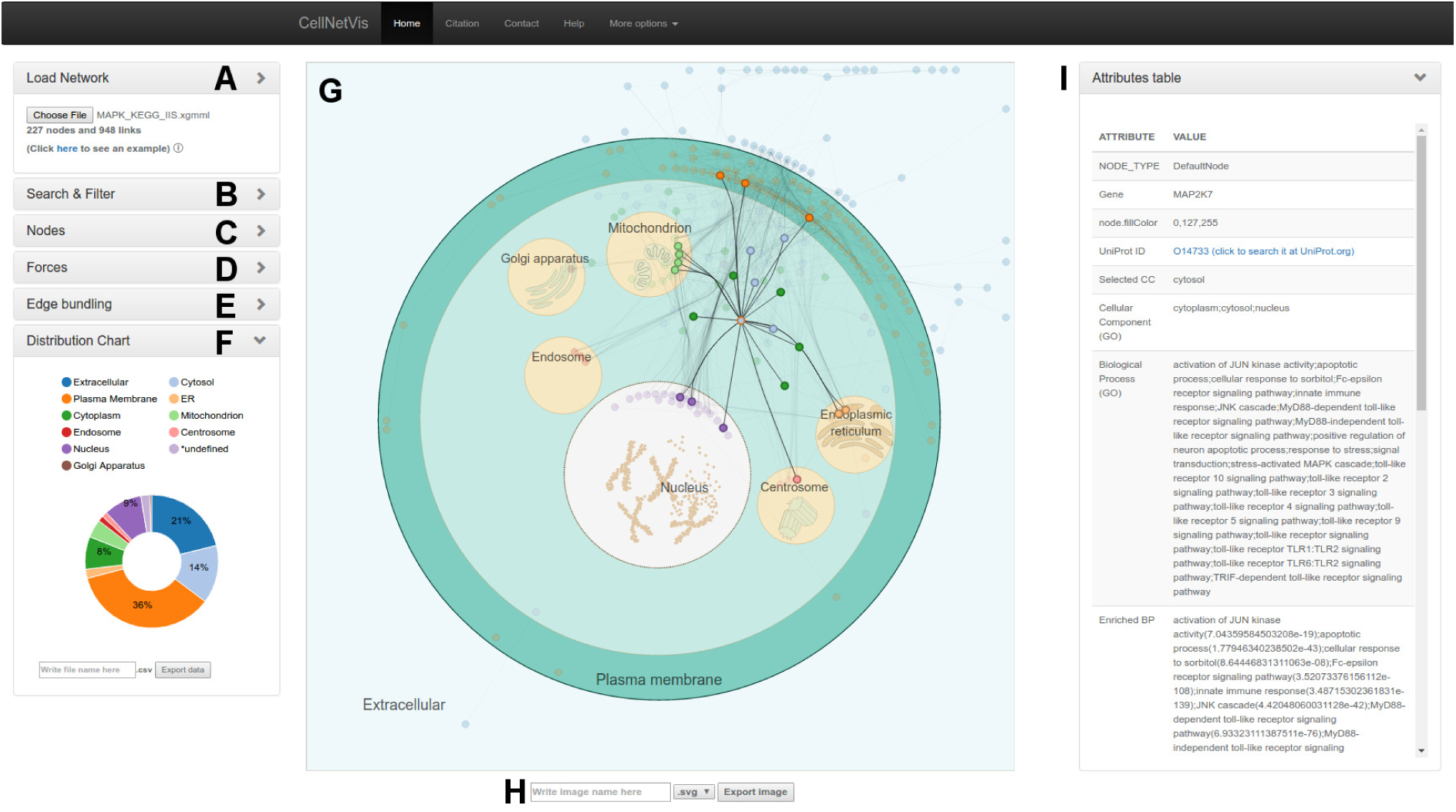
The CellNetVis interface. The main interface sections are indicated by lettering as follows: (A) Load network section, (B) Search section, (C) Nodes section, (D) Force-directed algorithm section, (E) Edge bundling section, (F) Donut chart section, (G) Cell diagram section, (H) Cell diagram export section, and (I) Node attributes table section.

Edge bundling may be applied to the network (Figures 2E e 2G). The effect is to group and smooth edges that flow along the same region of the display. Bundling edges typically reduces the visual density of the network layout, providing a clearer view of the relations among groups of nodes.

CellNetVis allows searching for nodes by label (Figure 2B). The tool then highlights the nodes matching the search. It is possible to hide unselected nodes using the filter functionality (“Filter” button). This allows users to focus the analysis in a fraction of the network. Node attributes may be displayed by our tool in a tabular fashion that includes a link to the UniProt website whenever the proper accession number is available (Figure 2I). Moreover, three network topology measures can be computed and added to each node: degree, betweenness, and clustering coefficient. The colors of the nodes can also be changed by using the drop-down list of nodes attributes (Figure 2C).

After loading a network, a donut chart showing the counts and percentages of nodes per cellular component is displayed on the bottom left side of the cell diagram (Figure 2F). Such chart may be exported in CSV, SVG and PNG formats. The cellular diagram may be exported as an SVG or PNG file.

The following sections describe two use cases and an additional comparison of CellNetVis against other available tools. In all cases, we used the same desktop computer with the following configuration: Chrome web browser version 56 (64-bit), Ubuntu 16.04 (64bit), Intel Core i7-2600K 3.4 GHz (launch date: 2011), GeForce GTX 750 Ti, and 8GB DDR3 RAM.

### Use Case 1: Comparison of GO and HPA subcellular compartments annotations on a *Homo sapiens* high-throughput network

We used 2097 proteins from the Human Protein Atlas [37] supportive data (Additional File 2) to construct a first neighbors network on IIS. A final large network containing 1942 nodes and 17498 links was then exported from IIS to CellNetVis to test the program capacity of handling large networks for a proper visualization and analysis (Figure 3A). Organelles were manually moved to improve the layout (Figure 3B) and edge bundling was turned on (Figure 3C). With these steps, the existence of edges and their frequency between cellular compartments became clearer. As expected, by comparing the donut chart information to the HPA data, the GO annotations ranking by the percentage of nodes distributed in each cellular component was similar to the HPA annotations ranking, particularly concerning the top (nucleus followed by cytoplasm, including cytoskeleton and cytosolic proteins) and bottom (microtubule organizing center) terms of the ranking (Additional File 3). This network is available through the CellNetVis Help page, and can be downloaded and uploaded or directly visualized at CellNetVis.

**Figure 3:**
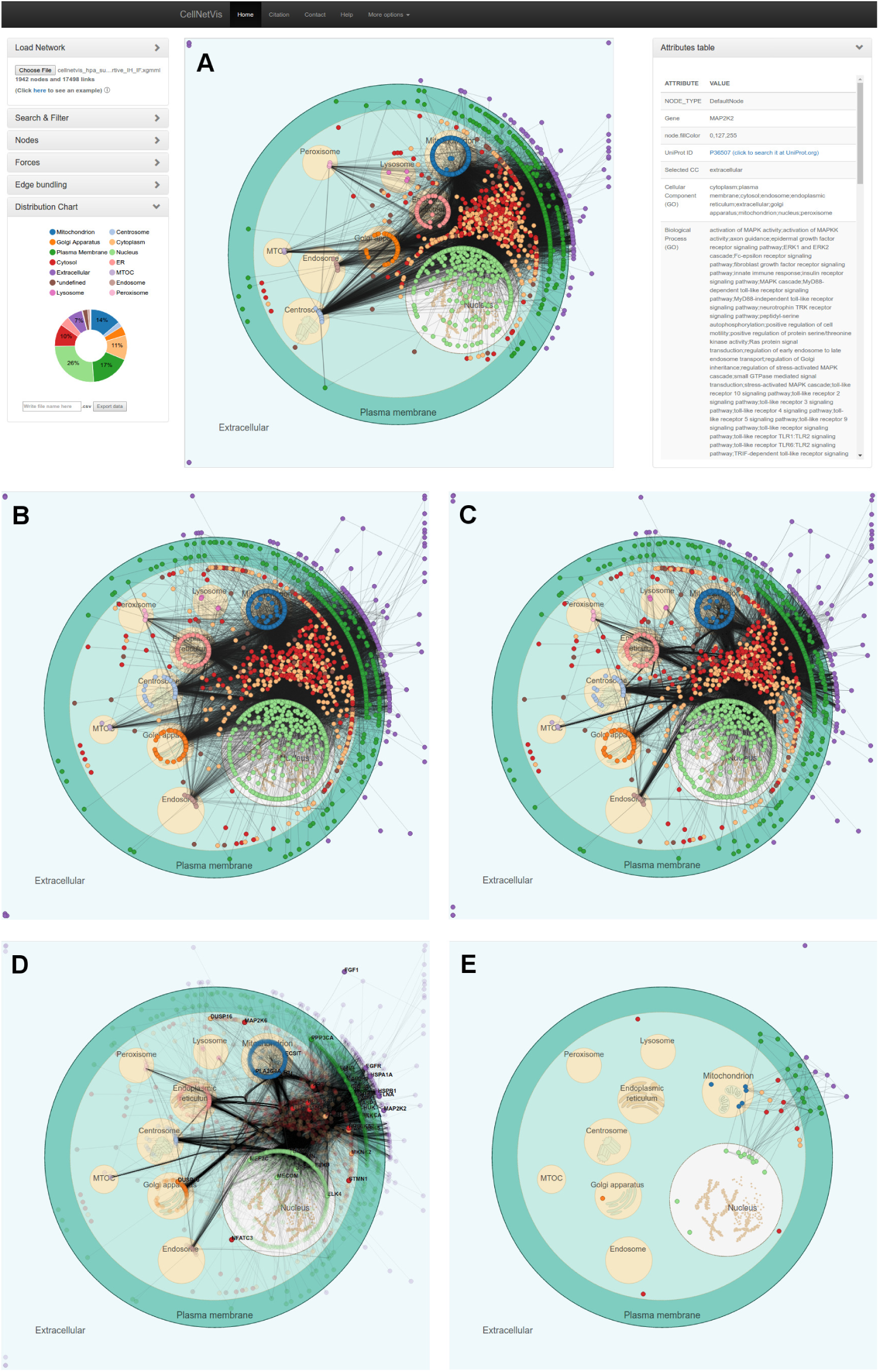
CellNetVis interface showing a human high-throughput network distributed on a cell diagram. (A) Visualization of the first neighbors network queried from IIS platform using 2097 proteins from the HPA supportive IH and IF data as input. The nodes’ colors were set to be displayed according to the “Selected CC” attribute. The MAP2K2 node was selected to show its attributes, as an example, on the table on the right side of the diagram. (B) Some organelles were moved and the force-directed algorithm was stopped. (C) Edge bundling was computed and displayed. (D) Proteins annotated to the human MAPK signaling pathway from KEGG database were searched in the network using the Find button and are shown as highlighted nodes. (E) Only the previously highlighted nodes corresponding to proteins annotated to the human MAPK signaling pathway are visible.

Besides being useful to connect the network to information regarding subcellular compartments, CellNetVis is also useful to analyze their interactions and pathways by setting node colors according to, e.g., the GO biological processes or KEGG [38] pathways, or by highlighting only the nodes annotated for a particular process/pathway, such as the MAPK signaling pathway (Additional File 4) depicted in Figure 3D. From the 257 proteins annotated as involved in the MAPK signaling pathway in the KEGG database (Additional File 4), only a fraction of them was found in the HPA network. Filtering enables this fraction of nodes to be visualized as a separate network, so that the user can more accurately analyze only the interactions pertinent to this specific pathway (Figure 3E). The force-directed algorithm may be restarted, and the layout computed considering only visible nodes.

CellNetVis handled 1942 nodes and 17498 edges, although still showing the hairball effect that most node-link approaches have. Despite the clutter, the user can see the distribution of nodes and edges in cellular components and has an overview of the network. Edge bundling also helps in the overview phase. The filtering function is important in exploration, as it allows the user to focus on areas and edges of interest while hiding everything else. The force-directed layout affects only visible nodes and the filtering function can be turned off at any time. Further techniques to change the visualization approach and reduce the hairball problem, e.g. Nodetrix [39] and Power Graphs [40], are scope for future work.

One limitation of CellNetVis is clear in this use case: although the system response was fast, the edge bundling took six minutes to complete and the nonoverlap functionality (repulsive force guided by *repulsive* value) had to be disabled. One alternative to the non-overlap functionality is to set a higher negative *charge* to nodes, which also has the effect of separating them. In our tests, Firefox browser loaded and showed the network three times faster than Chrome. Despite the good loading time, the system response on Chrome was much better than on Firefox. We tested the system response changing the network sizes (number of nodes and edges). According to our analysis, CellNetVis has a smaller and more stable response time on Chrome compared to Firefox (Additional File 5).

### Use Case 2: Visualization of the *Homo sapiens* MAPK signaling pathway organized in cellular compartments

We used 257 proteins from the human MAPK signaling pathway in the KEGG database (Additional File 4) to construct a first neighbors network on IIS. A final small network containing 227 nodes and 948 links was then exported from IIS to CellNetVis (Figure 4A). This file is also available on CellNetVis Help page to be downloaded and then uploaded or directly visualized at CellNetVis. Every time the user loads a different network, only the organelles corresponding to the GO cellular components annotations of that network are loaded in the cell diagram. Therefore, differently from the previously applied filter step on a larger network (Figure 3E), only the organelles annotated for the MAPK signaling pathway proteins are shown in this case. Due to the size of the network, the system response was good both on Chrome and Firefox, with Chrome still showing a larger speed.

**Figure 4:**
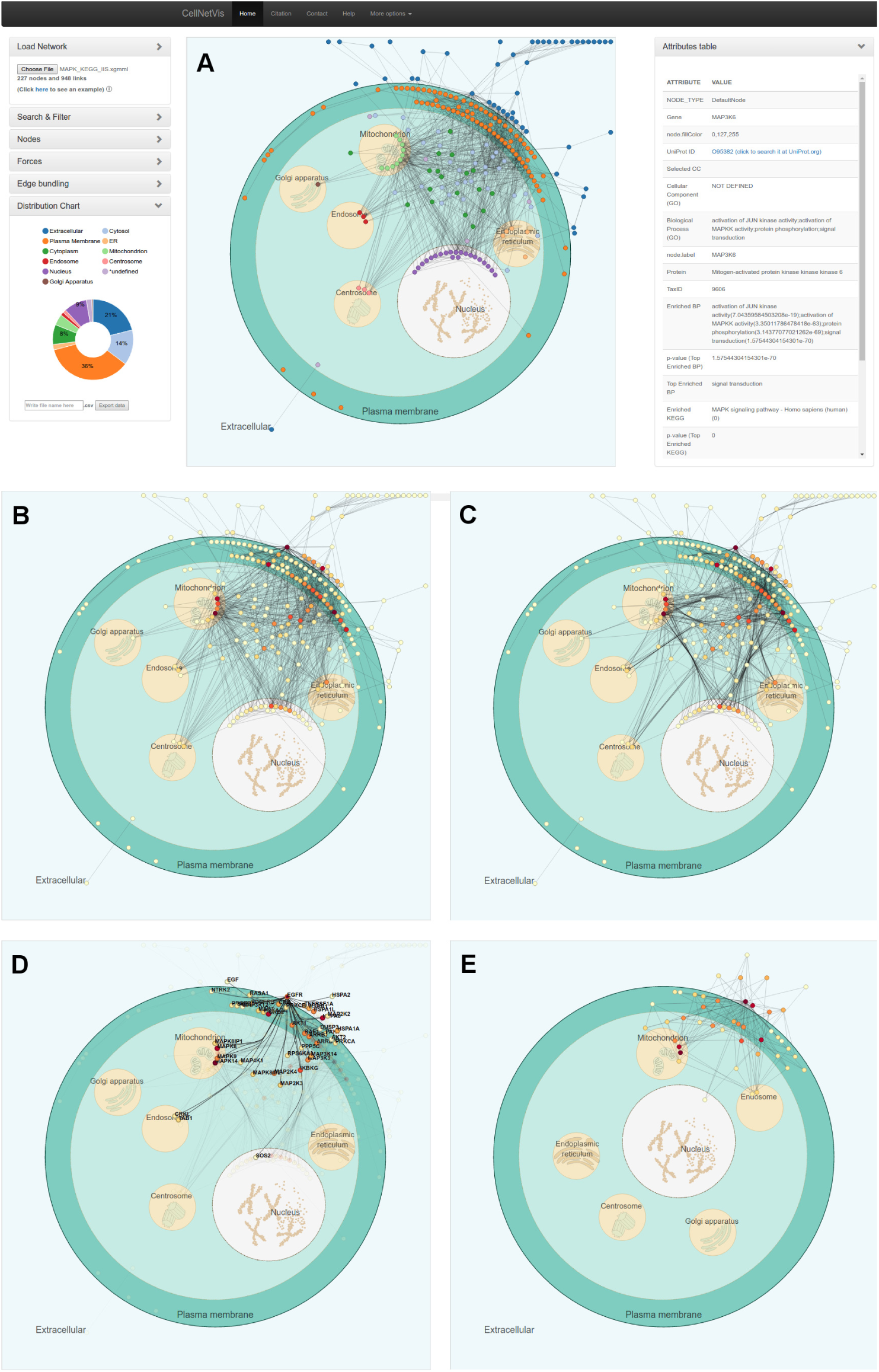
CellNetVis interface showing the human MAPK signaling pathway distributed in a cell diagram. (A) Visualization of the first neighbors network queried from IIS platform using 257 proteins annotated to the human MAPK signaling pathway from KEGG database as input. The nodes’ colors were set to be displayed according to the “Selected CC” attribute. The attributes of MAP3K6 are shown on the table on the right side of the diagram. (B) The nodes’ colors were set to be displayed according to the “[degree]” attribute. The force-directed algorithm was stopped. (C) Edge bundling wa computed and displayed. (D) The node with the darkest color, EGFR, was selected. The highlighted nodes correspond to the EGFR’s first neighbors, after EGFR’s node selection. (E) Only highlighted nodes are visible and force-directed layout was restarted. Organelles were moved to improve the layout.

The nodes were colored by their degree, in order to show the hubs (nodes with the highest connectivity), representing the proteins responsible for the major signal integration and transduction in the pathway (Figure 4B). Edge bundling was applied for a better visualization of the main paths of signal flow in the network (Figure 4C). From this analysis, we observe that the main paths occur between the extracellular region and plasma membrane, between the plasma membrane and mitochondrion, endoplasmic reticulum, endosome, centrosome or nucleus, and between the cytosol and the previously mentioned organelles. We can also observe that the hubs (dark red) are mainly located in the extracellular region, plasma membrane, and mitochondrion.

By clicking on the node with the darkest color (the highest degree), its label appears (EGFR), the table is updated to show EGFR node attributes on the right side of the diagram, and only the first neighbors of EGFR are highlighted in the network (Figure 4D). This analysis showed that EGFR interacts with proteins on the extracellular region, plasma membrane, cytosol, mitochondrion, endosome, and nucleus. By looking at the “Cellular Component (GO)” line on the nodes attributes table, we observe that EGFR is not annotated to localize at mitochondria. This suggests that EGFR may interact with those mitochondrial proteins at other subcellular compartments where they also exist, such as the case of MAPK14, which interaction may occur in the cytoplasm or nucleus. In Figure 4E, organelles were moved and the force-directed layout restarted to create a layout that focuses on the subnetwork topology instead of on concentration and flow of interactions through the cell compartments.

### Comparison with available tools

A comparison was performed between the force-directed layout of CellNetVis, the multiple force-directed layout of Mosaic [8] plugin, and the grid layout of Cerebral [7] plugin and CerebralWeb [23]. Although Cell Illustrator Online (CIO) [25] is capable of showing networks inside a cell diagram, the modeling and cell diagram must be manually set up, the tool focuses on the molecular mechanisms and is not freely available, thus, it was not considered in the comparison.

Our focus is freely available systems that can automatically partition the network into a cellular diagram and display a simple and interactive overview in a fast and easy way. Although Cerebral and CerebralWeb do not display a cell diagram, they can automatically separate the network into partitions. Also, CerebralWeb is freely available and can be integrated into web systems. Mosaic is not web-based, but it can automatically place nodes over a cell diagram, therefore it was also considered in the comparison. The main characteristics in contrast with CellNetVis are detailed in Table 2

**Table 2:**
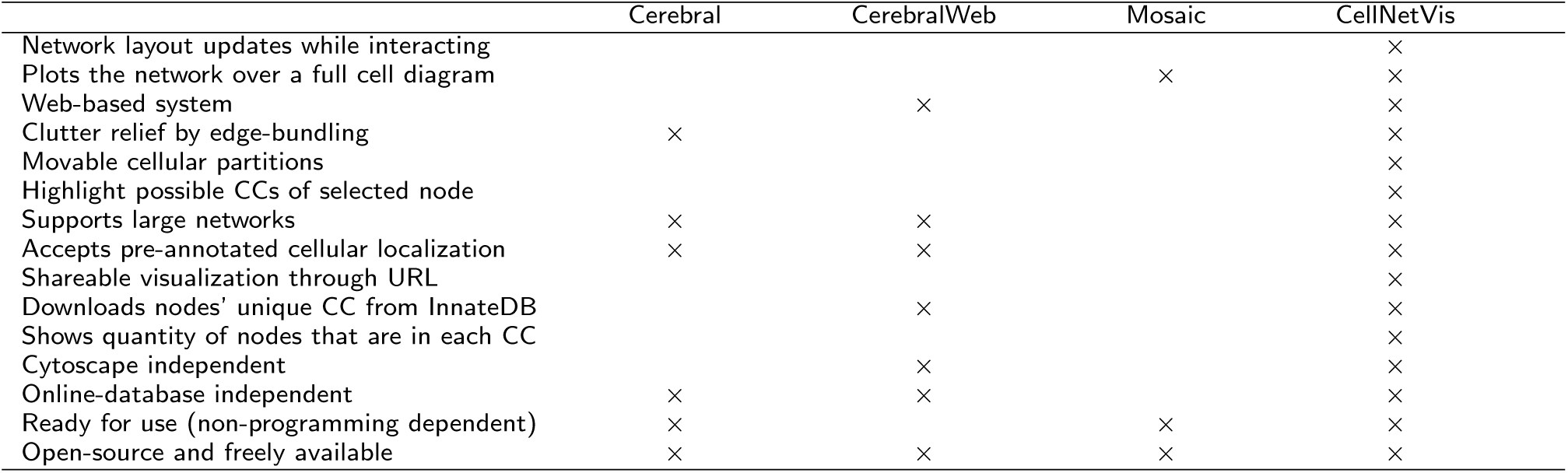
Carachteristics of CellNetVis, Cerebral, CerebralWeb, and Mosaic.

Mosaic is a Cytoscape (desktop) plugin which partitions a network into subnetworks based on GO Biological Process annotation. Each subnetwork is shown in a different cellular diagram. If a node has more than one value to this attribute, the node is duplicated. Since the tool uses the force-directed algorithm to place nodes, the layout is similar to CellNetVis. However, the system was designed to load the small subnetworks created based on nodes annotations. Overlap of nodes is very common even for small networks.

We could not replicate the results described in [8] using Mosaic since it is out of date and could not download its required databases. Therefore we decided to create an analysis based on the Yeast example network available at Mosaic web page (http://nrnb.org/tools/mosaic/). We created a new annotated network (642 nodes and 7785 edges) with all interactions, found by the IIS, between all the listed genes from the Yeast example. Then, we visualized on CellNetVis the network (Additional File 6A and 6B) and subnetworks created by filtering the biological process annotations: ‘regulation of transcription’, ‘metabolic process’, ‘golgi to vacuole transport’, and ‘intracellular protein transport’ (Additional File 6C, 6D, 6E and 6F, respectively). Using as basis the figures ^[1][2][3]^ displayed on Mosaic web page, section *Navigating the results*, CellNetVis performed better, since nodes did not overlap on any of the displayed subnetworks and their topology was clear.

Regarding the Cerebral plugin and CerebralWeb, the network layout algorithm is modeled after hand-drawn pathway diagrams, where nodes are restricted to a regular lattice grid that provides room for labels and eliminates overlapping nodes [7]. The main difference to CellNetVis is the use of a grid layout to position nodes on horizontal layers, one over the other, so as to resemble subcellular compartments. However, the use of horizontal layers for this purpose restricts cell layers to the five major subcellular compartments, which are positioned by Cerebral from top to bottom in the following order: extracellular, cell surface, plasma membrane, cytoplasm, and nucleus. For instance, the majority of organelles, which are naturally localized in the cytoplasm, cannot be drawn inside the cytoplasm layer in Cerebral, only as horizontal layers on the top, bottom or between the other ones (e.g. below nucleus, as default), which is not consistent with an appropriate cellular view (Additional File 7). The same happens in the web-based version of the system.

Comparing the loading and drawing times for a large network composed of 1942 nodes, Cerebral took about 4 minutes, while CellNetVis took half the time to load the network file, to check for duplicate nodes and edges, to create the data structure, to start the force-directed layout, and nearly stabilize the force system and to display a consistent layout of the network topology. For a small network composed of 227 nodes, Cerebral took 10 seconds, while CellNetVis took approximately 1 second.

To compare the layout created by CerebralWeb and CellNetVis we created the displays for the networks from Use Case 1 (Additional File 7A vs. Figure 3A) and Use Case 2 (Additional File 7B vs. Figure 4A). In both cases, CerebralWeb was not capable of clearly representing the density of interaction between compartments as CellNetVis does. For instance, in Figure 3A we can see that there are more interactions between mitochondrion and nucleus than between endoplasmic reticulum and nucleus; in CerebralWeb it is not possible to see this pattern (Additional File 7B). Moving the organelles on CellNetVis also allows the user to check this type of information. Considering the overview of the network on CerebralWeb (Additional File 7B), the only information we can visually identify in the diagram is the distribution of nodes over the compartments. This information can be more easily identified in CellNetVis through the distribution chart (Figure 2F). Thus the overview created by CellNetVis is more informative than the one created by CerebralWeb. In contrast to Cerebral, CerebralWeb can draw large networks fast, but the layout is not as good as the layout computed by the Cerebral plugin (Additional File 7A and 7C). We integrated CerebralWeb to CellNetVis system, which can be accessed through the “More options” top-menu item after loading a network. Both CerebralWeb and CellNetVis layout were displayed almost instantly after loading the network file from Use Case 2.

Another advantage of CellNetVis concerns the highlight and filtering of nodes or pathways in a complex network. As shown in Figure 3A, when a network is large there are many nodes overlapping. CellNetVis allows the user to filter nodes based on a search query. These filtered nodes can be automatically repositioned. This functionality and interactivity improves the network display and exploration and is not possible in Cerebral, where the layout is pre-calculated. Cerebral only allows the highlight of neighbors for a selected node and is able to recalculate the layout as a second drawing step, but only considering all the nodes. The web version needs to be programmed to be used with these features, despite being implemented as a module of the CerebralWeb Javascript library.

One fact that could be considered a limitation of CellNetVis appears in Figure 3A, where nodes overlap at a high rate due to network size. However, the overlap of nodes is what allows the density of edges between organelles clear supporting the overview task and being more informative than the non-overlap layout created by CerebralWeb algorithm. CellNetVis can show at the overview step the connectivity among compartments (edges densities), the distribution of nodes (chart distribution), and give details according to the user interactions by search, filtering, and selection of nodes. After filtering a large network, for instance, the charges of nodes or *repulsive* value can be increased to drastically reduce overlapping effect. Considering the critical execution time that happens on general web-applications, we could say for both web-based layouts compared in this section, that CellNetVis and CerebralWeb focus on being fast enough to be used with considerably large networks. CellNetVis lets nodes overlap at a high rate when networks are large, but keeps the dynamic aspect of the layout and accentuate the concentration of edges (Figure 3), whereas CerebralWeb layout algorithm avoids the overlap of nodes but is not dynamic and hides the overview of the network topology (Additional File 7B).

The positioning algorithm of CellNetVis works well with both small and large networks and supports more directly the visualization pipeline described by Schneiderman [41]: overview, followed by zoom and filter, then details on demand. If a user modifies the position of a node or organelle in the network representation, CellNetVis is able to recalculate the position of the other nodes instantly, while Cerebral and CerebralWeb are not. Moving a node or organelle can highlight certain aspects of the data (Additional File 8A). For instance, if nodes are too close inside the cell, the user can separate nodes to let the topology clear (Additional File 8B). This cannot be accomplished by using CerebralWeb (Additional File 8C) or Cerebral (Additional File 8D). CellNetVis was shown to be a more flexible tool throughout user interaction tasks. Due to characteristics of the layout algorithm, the movement of a node in CellNetVis is not so smooth and precise, but still usable and useful.

## Conclusions

CellNetVis is a free and open-source web-based software for displaying biomolecular networks in a cell diagram. It is capable of displaying complex information related to networks, nodes and edges, as well as their relations with cell partitions. While being better suited for small and medium-sized networks, CellNetVis is also capable of handling large networks. In comparison with other algorithms and tools, CellNetVis has shown to be competitive, particularly for a dynamic exploration of complex networks over a consistent representation of a full cell on the Web. CellNetVis is being used by the IIS as its main visualization system. CellNetVis may also be coupled with different anottation softwares using the XGMML format to exchange data, providing an interesting analysis layer.

## AVAILABILITY AND REQUIREMENTS

**Project name**: CellNetVis

**Project home page**: http://www.lge.ibi.unicamp.br/cellnetvis

**Operating system(s)**: Platform independent

**Programming language**: JavaScript and HTML

**Other requirements**: Web browsers Chrome 46+, Firefox 40+, or IE 10+

**License**: GPLv3

### Author’s contributions

The system was proposed by GVM. HH and GVM designed the system that was implemented by HH and tested by GVM, MFC, RM and GPT. GVM proposed the use cases and conducted them with HH. GVM and RM supervised the project. All authors wrote and approved the final manuscript.

## Acknowledgements

The authors acknowledge Hugo Henrique Slepicka for the technical support related to IIS.

## List of abbreviations used

IIS: *Integrated Interactome System*
GO: *Gene Ontology*
HPA: *Human Protein Atlas*
IH: *ImmunoHistochemistry*
IF: *ImmunoFluorescence*
Selected CC: *Selected Cellular Component*
CC: *Cellular Component*
MAPK: *Mitogen-Activated Protein Kinases*
EGFR: *Epidermal Growth Factor Receptor*
SBGN: *Systems Biology Graphical Notations*
SVG: *Scalable Vector Graphics*
PNG: *Portable Network Graphics*
XGMML: *eXtensible Graph Markup and Modeling Language*
URL: *Uniform Resource Locator*
CIO: *Cell Illustrator Online*.

## Additional Files

Additional File 1 — Diagram of the force-directed layout constraint algorithm

The diagram represents the basic concept about how nodes’ positions are redefined by our constraining algorithm during the force-direct layout iterations. It shows how a node is moved from cytosol to the inner-border of an organelle defined in its Selected CC attribute (A) and how a node that should be in the “cytosol” is moved from an organelle to its outer-border (B).

Additional File 2 — Supportive immunohistochemistry (IH) and immunofluorescence (IF) data from the Human Protein Atlas (http://www.proteinatlas.org/about/download) The table contains a subset of the data in the Human Protein Atlas version 13 corresponding to the data downloaded in TAB format, filtered by the supportive IH and IF information.

Additional File 3 — HPA and GO subcellular compartments annotations Tables showing the HPA and GO subcellular compartments annotations ranked by the absolute number and percentage of proteins assigned to each term.

Additional File 4 — Gene annotations retrieved from the *Homo sapiens* MAPK signaling pathway of the KEGG database.

The gene symbols were used as queries in the Search box of CellNetVis to highlight the proteins involved in this pathway.

Additional File 5 — Tests of CellNetVis’s response time on Firefox and Chrome

The table shows how fast was the interaction with the graphic user interface (GUI) and the constrained force-directed algorithm when executed on Firefox and Chrome on Ubuntu/Linux system.

Additional File 6 — Visualization of Yeast subnetworks filtered by specific biological processes

(A) and (B) represent the complete Yeast network formed by 642 nodes and 7785 edges. The network was filtered according to the following biological processes: ‘regulation of transcription’ (C), ‘metabolic process’ (D), ‘golgi to vacuole transport’ (E), and ‘intracellular protein transport’ (F).

Additional File 7 — Visualization of a large and of a small network on the Cerebral Cytoscape plugin (A and C) and on CerebralWeb (B and D).

(A and B) Large network generated from the HPA supportive data. The drawing took approximately 3.5 min. in Cerebral (A) and 6 s. in CerebralWeb (B). (C and D) Small network generated from the human MAPK signaling pathway from KEGG database. The drawing took approximately 5 s. on Cerebral (C) and 1 s. on (D). HPA: Human Protein Atlas; MAPK: Mitogen-activated protein kinases.

Additional File 8 — Visualization of the RIG-I-like receptor signaling pathway

Visualization of the network formed by the interactions within the “RIG-I-like receptor signalling pathway (KEGG)” in *Homo sapiens*, downloaded from InnateDB (http://innatedb.ca/interactionSearch.do?from=pw&exPathwayXref=&pathwayFilter=5713&pathwayXrefDB=&pathwayXref=&listType=interaction&coreInteractors=true) as a XGMML file and loaded on CellNetVis, CerebralWeb and Cerebral. (A and B) Visualization of the network on CellNetVis before (A) and after (B) manually separating nodes with high degree (dark red). The same network was draw on CerebralWeb (C) and Cerebral (D) for comparison. CellNetVis was shown to be a more flexible tool through user interaction.

## Footnotes

1 http://nrnb:org/tools/mosaic/images/mosaicresults:png

2 http://nrnb:org/tools/mosaic/images/mosaic-subnetwork:png

3 http://nrnb:org/tools/mosaic/images/mosaic-selectnodes:png

## References

1. Ellson, J., Gansner, E., Koutsofios, L., North, S.C., Woodhull, G.: In: Mutzel, P., Jünger, M., Leipert, S. (eds.) Graphviz—Open Source Graph Drawing Tools, pp. 483–484. Springer, Berlin, Heidelberg (2002). doi:10.1007/3-540-45848-457

2. Bastian, M., Heymann, S., Jacomy, M.: Gephi: An Open Source Software for Exploring and Manipulating Networks. In: ICWSM (2009). http://www.aaai.org/ocs/index.php/ICWSM/09/paper/view/154

3. V. Batagelj, a.M.: Pajek – program for large network analysis. Connections, 47–57 (1998). doi:10.1.1.27.9156

4. Martins, R.M., Andery, G.F., Heberle, H., Paulovich, F.V., Andrade Lopes, A., Pedrini, H., Minghim, R.: Multidimensional Projections for Visual Analysis of Social Networks. Journal of Computer Science and Technology 27(4), 791–810 (2012). doi:10.1007/s11390-012-1265-5

5. Shannon, P., Markiel, A., Ozier, O., Baliga, N.S., Wang, J.T., Ramage, D., Amin, N., Schwikowski, B., Ideker, T.: Cytoscape: a software environment for integrated models of biomolecular interaction networks. Genome research 13(11), 2498–504 (2003). doi:10.1101/gr.1239303

6. Auber, D., Archambault, D., Bourqui, R., Delest, M., Dubois, J., Pinaud, B., Lambert, A., Mary, P., Mathiaut, M., Melancon, G.: Tulip III. In: Encyclopedia of Social Network Analysis and Mining, (2014). doi:10.1007/978-1-4614-6170-8315. https://hal.archives-ouvertes.fr/hal-01096759

7. Barsky, A., Gardy, J.L., Hancock, R.E.W., Munzner, T.: Cerebral: a Cytoscape plugin for layout of and interaction with biological networks using subcellular localization annotation. Bioinformatics 23(8), 1040–1042 (2007). doi:10.1093/bioinformatics/btm057

8. Zhang, C., Hanspers, K., Kuchinsky, A., Salomonis, N., Xu, D., Pico, A.R.: Mosaic: making biological sense of complex networks. Bioinformatics 28(14), 1943–1944 (2012). doi:10.1093/bioinformatics/bts278

9. Paduano, F., Forbes, A.: Extended LineSets: a visualization technique for the interactive inspection of biological pathways. BMC 9(Suppl 6), 4 (2015). doi:10.1186/1753-6561-9-S6-S4

10. Lex, A., Partl, C., Kalkofen, D., Streit, M., Gratzl, S., Wassermann, A.M., Schmalstieg, D., Pfister, H.: Entourage: Visualizing Relationships between Biological Pathways using Contextual Subsets. IEEE Transactions on Visualization and Computer Graphics 19(12), 2536–2545 (2013). doi:10.1109/TVCG.2013.154

11. Dang, T., Murray, P., Aurisano, J., Forbes, A.: ReactionFlow: an interactive visualization tool for causality analysis in biological pathways. BMC Proceedings 9(Suppl 6), 6 (2015). doi:10.1186/1753-6561-9-S6-S6

12. Garcia, O., Saveanu, C., Cline, M., Fromont-Racine, M., Jacquier, A., Schwikowski, B., Aittokallio, T.: GOlorize: a Cytoscape plug-in for network visualization with Gene Ontology-based layout and coloring. Bioinformatics 23(3), 394–396 (2007). doi:10.1093/bioinformatics/btl605

13. Dwyer, T.: Scalable, Versatile and Simple Constrained Graph Layout. Computer Graphics Forum 28(3), 991–998 (2009). doi:10.1111/j.1467-8659.2009.01449.x

14. Dwyer, T., Robertson, G.: Layout with Circular and Other Non-linear Constraints Using Procrustes Projection. In: Graph Drawing: 17th International Symposium, GD 2009, Chicago, IL, USA, September 22–25, 2009. Revised Papers, pp. 393–404 (2010). doi:10.1007/978-3-642-11805-0 37. http://link.springer.com/10.1007/978-3-642-11805-0 37

15. Didimo, W., Montecchiani, F.: Fast layout computation of clustered networks: Algorithmic advances and experimental analysis. Information Sciences 260(1), 185–199 (2014). doi:10.1016/j.ins.2013.09.048

16. Didimo, W., Montecchiani, F.: Fast Layout Computation of Hierarchically Clustered Networks: Algorithmic Advances and Experimental Analysis. 2012 16th International Conference on Information Visualisation, 18–23 (2012). doi:10.1109/IV.2012.14

17. Schuhmacher, A.: Software Visualization via Hierarchic Graphs. PhD thesis, Karlsruhe Institute of Technology (2015)

18. Baur, M., Brandes, U.: Multi-circular layout of micro/macro graphs. Lecture Notes in Computer Science (including subseries Lecture Notes in Artificial Intelligence and Lecture Notes in Bioinformatics) 4875 LNCS, 255–267 (2008). doi:10.1007/978-3-540-77537-926

19. Dogrusoz, U., Giral, E., Cetintas, A., Civril, A., Demir, E.: A layout algorithm for undirected compound graphs. Information Sciences 179(7), 980–994 (2009). doi:10.1016/j.ins.2008.11.017

20. Archambault, D., Munzner, T., Auber, D.: Tugging graphs faster: Efficiently modifying path-preserving hierarchies for browsing paths. IEEE Transactions on Visualization and Computer Graphics 17(3), 276–289 (2011). doi:10.1109/TVCG.2010.60

21. Altarawneh, R., Schultz, J., Humayoun, S.R.: CluE: An algorithm for expanding clustered graphs. IEEE Pacific Visualization Symposium, 233–237 (2014). doi:10.1109/PacificVis.2014.18

22. Krause, F., Schulz, M., Ripkens, B., Flottmann, M., Krantz, M., Klipp, E., Handorf, T.: Biographer: web-based editing and rendering of SBGN compliant biochemical networks. Bioinformatics 29(11), 1467–1468 (2013). doi:10.1093/bioinformatics/btt159

23. Frias, S., Bryan, K., Brinkman, F.S.L., Lynn, D.J.: CerebralWeb: A cytoscape.js plug-in to visualize networks stratified by subcellular localization. Database 2015, 1–4 (2015). doi:10.1093/database/bav041

24. Kojima, K., Nagasaki, M., Miyano, S.: Fast grid layout algorithm for biological networks with sweep calculation. Bioinformatics 24(12), 1433–1441 (2008). doi:10.1093/bioinformatics/btn196

25. Nagasaki, M., Saito, A., Jeong, E., Li, C., Kojima, K., Ikeda, E., Miyano, S.: Cell illustrator 4.0: a computational platform for systems biology. In silico biology 10(1, 2), 5–26 (2010). doi:10.3233/978-1-60750-704-8-160

26. Ashburner, M., Ball, C.A., Blake, J.A., Botstein, D., Butler, H., Cherry, J.M., Davis, A.P., Dolinski, K., Dwight, S.S., Eppig, J.T., Harris, M.A., Hill, D.P., Issel-Tarver, L., Kasarskis, A., Lewis, S., Matese, J.C., Richardson, J.E., Ringwald, M., Rubin, G.M., Sherlock, G.: Gene ontology: tool for the unification of biology. The Gene Ontology Consortium. Nature genetics 25(1), 25–9 (2000). doi:10.1038/75556

27. Punin, J., Krishnamoorthy, M.: XGMML (eXtensible Graph Markup and Modeling Language) 1.0 Draft Specification. (2001)

28. Carazzolle, M.F., De Carvalho, L.M., Slepicka, H.H., Vidal, R.O., Pereira, G.A.G., Kobarg, J., Meirelles, G.V.: IIS - Integrated Interactome System: A web-based platform for the annotation, analysis and visualization of protein-metabolite-gene-drug interactions by integrating a variety of data sources and tools. PLoS ONE 9(6), 100385 (2014). doi:10.1371/journal.pone.0100385

29. Breuer, K., Foroushani, A.K., Laird, M.R., Chen, C., Sribnaia, A., Lo, R., Winsor, G.L., Hancock, R.E.W., Brinkman, F.S.L., Lynn, D.J.: Innatedb: systems biology of innate immunity and beyond—recent updates and continuing curation. Nucleic Acids Research 41(D1), 1228 (2013). doi:10.1093/nar/gks1147

30. Yates, A., Akanni, W., Amode, M.R., Barrell, D., Billis, K., Carvalho-Silva, D., Cummins, C., Clapham, P., Fitzgerald, S., Gil, L., Girón, C.G., Gordon, L., Hourlier, T., Hunt, S.E., Janacek, S.H., Johnson, N., Juettemann, T., Keenan, S., Lavidas, I., Martin, F.J., Maurel, T., McLaren, W., Murphy, D.N., Nag, R., Nuhn, M., Parker, A., Patricio, M., Pignatelli, M., Rahtz, M., Riat, H.S., Sheppard, D., Taylor, K., Thormann, A., Vullo, A., Wilder, S.P., Zadissa, A., Birney, E., Harrow, J., Muffato, M., Perry, E., Ruffier, M., Spudich, G., Trevanion, S.J., Cunningham, F., Aken, B.L., Zerbino, D.R., Flicek, P.: Ensembl 2016. Nucleic Acids Research 44(D1), 710 (2015). doi:10.1093/nar/gkv1157

31. McEntyre, J.: Linking up with entrez. Trends in Genetics 14(1), 39–40 (1998). doi:10.1016/S0168-9525(97)01325-5

32. Magrane, M., Consortium, U.: UniProt Knowledgebase: a hub of integrated protein data. Database: the journal of biological databases and curation 2011, 009 (2011). doi:10.1093/database/bar009

33. Harrower, M., Brewer, C.A.: ColorBrewer.org: An Online Tool for Selecting Colour Schemes for Maps. The Cartographic Journal 40(1), 27–37 (2003). doi:10.1179/000870403235002042

34. Bostock, M., Ogievetsky, V., Heer, J.: D3: Data-Driven Documents. IEEE transactions on visualization and computer graphics 17(12), 2301–9 (2011). doi:10.1109/TVCG.2011.185

35. Samet, H.: The quadtree and related hierarchical data structures. ACM Comput. Surv. 16(2), 187–260 (1984). doi:10.1145/356924.356930

36. Holten, D., van Wijk, J.J.: Force-Directed Edge Bundling for Graph Visualization. Computer Graphics Forum 28(3), 983–990 (2009). doi:10.1111/j.1467-8659.2009.01450.x

37. Uhlen, M., Oksvold, P., Fagerberg, L., Lundberg, E., Jonasson, K., Forsberg, M., Zwahlen, M., Kampf, C., Wester, K., Hober, S., Wernerus, H., Björling, L., Ponten, F.: Towards a knowledge-based Human Protein Atlas. Nature biotechnology 28(12), 1248–50 (2010). doi:10.1038/nbt1210-1248

38. Kanehisa, M., Goto, S., Kawashima, S., Nakaya, A.: The KEGG databases at GenomeNet. Nucleic acids research 30(1), 42–6 (2002)

39. Henry, N., Fekete, J.-D., McGuffin, M.J.: NodeTrix: a Hybrid Visualization of Social Networks. IEEE Transactions on Visualization and Computer Graphics 13(6), 1302–1309 (2007). doi:10.1109/TVCG.2007.70582

40. Wang, Y., Thilmony, R., Gu, Y.Q.: NetVenn: an integrated network analysis web platform for gene lists. Nucleic Acids Research 42(W1), 161–166 (2014). doi:10.1093/nar/gku331

41. Shneiderman, B.: The eyes have it: a task by data type taxonomy for information visualizations. In: Proceedings 1996 IEEE Symposium on Visual Languages, pp.336–343. IEEE Comput. Soc. Press. doi:10.1109/VL.1996.545307

